# New Viral Biogeochemical Roles Revealed Through Metagenomic Analysis of Lake Baikal

**DOI:** 10.1101/2020.04.02.019802

**Authors:** FH Coutinho, PJ Cabello-Yeves, R Gonzalez-Serrano, R Rosselli, M López-Pérez, TI Zemskaya, AS Zakharenko, VG Ivanov, F Rodriguez-Valera

## Abstract

Lake Baikal is the largest body of liquid freshwater on Earth. Previous studies have described the microbial composition of this habitat but the viral communities from this ecosystem have not been characterized in detail. Here we describe the viral diversity of this habitat across depth and seasonal gradients. We discovered 19,475 *bona fide* viral sequences, which are derived from viruses predicted to infect abundant and ecologically important taxa that reside in Lake Baikal, such as Nitrospirota, Methylophilaceae and Crenarchaeota. Diversity analysis revealed significant changes in viral community composition between epipelagic and bathypelagic zones. Analysis of the gene content of individual viral populations allowed us to describe one of the first bacteriophages that infect Nitrospirota, and their extensive repertoire of auxiliary metabolic genes that might enhance carbon fixation through the reductive TCA cycle. We also described bacteriophages of methylotrophic bacteria with the potential to enhance methanol oxidation and the S-adenosyl-L-methionine cycle. These findings unraveled new ways by which viruses influence the carbon cycle in freshwater ecosystems, namely by using auxiliary metabolic genes that act upon metabolisms of dark carbon fixation and methylotrophy. Therefore, our results shed light on the processes through which viruses can impact biogeochemical cycles of major ecological relevance.

## Introduction

Lake Baikal is the largest and deepest lake on Earth (1,2)□. Its uniqueness also lies in its extreme oligotrophy, ice-covered periods of up to 4-4.5 months per year, and an oxic water column throughout all depths (3)□. The lake is permanently mixed and only undergoes stratification for a brief period of time during summer in its first 100 meters (4)□. The surface of Lake Baikal freezes during winter, so that below the ice layer water temperatures approach 0°C while towards deeper waters temperature raises slightly to a maximum of 4°C. In summer, the ice layer melts and surface water temperature raises to nearly 12°C, only to decrease rapidly towards deeper waters (below 50m) to the same 4°C that are kept all year around for the deep water mass. Recent metagenomic studies have analysed the microbiome of sub-ice epipelagic and bathypelagic waters, revealing the key microbes that dwell at this ecosystem as well as the ecological processes in which they are involved (5,6)□. These studies have shed light on the taxonomic composition of the Lake Baikal microbiome and the contributions of these microbes to biogeochemical cycles. Nevertheless, one important component of this ecosystem has not yet been characterized in detail: the viruses.

Viruses play key roles in the functioning of aquatic ecosystems (7,8)□. They mediate recycling of organic matter in these habitats by lysing host cells, which leads to the daily release of billions of tons of organic carbon (9)□. Yet the influence of viruses over aquatic microbiomes is not limited to killing. They can also modify host metabolism during infection through the expression of auxiliary metabolic genes (AMGs), which redirect host metabolism towards pathways that promote the production of viral particles, such as the nucleotide metabolism, which yields the building blocks necessary to synthesize the genomes of the viral progeny. (10,11)□. There are multiple mechanisms by which viruses make use of AMGs to re-direct host metabolism. Among the noteworthy examples of AMGs are included genes of photosynthesis and carbon fixation in viruses of Cyanobacteria (12)□, genes for sulfur oxidation in viruses of Proteobacteria (13)□, and genes for carbon metabolism, phosphorus metabolism, and protein synthesis spread across multiple host taxa (14–16)□, The prevalence and diversity of AMGs varies across ecosystems (17,18)□, hence the repertoire of molecular functions encoded by AMGs is expected to change in response to the metabolic constraints faced by their hosts across different ecosystems as well.

Extensive research has been conducted on the biodiversity, ecology and AMGs of viruses from marine ecosystems (15.19). Meanwhile, viruses from freshwater ecosystems have received much less attention. Consequently, little is known about the environmental drivers of community composition, biodiversity, and auxiliary metabolic genes in these ecosystems. The use of metagenomics has made it possible to describe viruses that infect the most representative freshwater microbes, such as acI Actinobacteria (20,21)□ and SAR11 (22,23)□. Studies focused on Czech reservoirs and Lake Biwa (Japan) have reported on the extensive effects of stratification on freshwater viral communities, host prevalence and on their key roles as recyclers of organic matter (21,24)□. Specifically, these studies found that in these stratified lakes, there is constantly shifting viral community in the epilimnion and a more stable community that dwells in the hypolimnion. Meanwhile, in Lake Baikal, recent studies have described a phage putatively infecting *Polynucleobacter sp*. (5)□, the virome associated with diseased sponges (25)□, and viromes from the first epipelagic zone which suggested that the Baikal virome undergoes changes in composition across seasons (26)□

Here we sought to perform an in-depth characterization of the viral communities from Lake Baikal. Our sampling strategy included retrieving samples at the epipelagic (photic), mesopelagic (aphotic) and bathypelagic (aphotic) zones during winter and summer. In total, ten cellular metagenomes were obtained from which viral sequences were identified. We used computational methods to assign putative hosts to viral sequences and to classify them taxonomically. With that information we investigated shifts in community composition regarding taxonomic affiliation and target hosts that were driven by depth and season. Next we described in detail the gene content of novel viruses infecting some of the most abundant members of the Lake Baikal microbiome and their repertoire of auxiliary metabolic genes that include key metabolic processes that have not been described before in other viruses.

## Results and Discussion

### Depth variations of Archaeal and Bacterial communities in Lake Baikal

We have analysed a total of ten metagenomes sequenced from different habitats of Lake Baikal. The four datasets that represented winter microbiomes from sub-ice samples have previously been described in detail (5,6)□. We expanded this set of samples by generating a new set of six metagenomes obtained from similar geographical coordinates but collected during summer. These included two epipelagic samples (photic, 5m and 20m), two mesopelagic samples (aphotic, 300m and 390m) at a site near methane seep that is notorious for high methane concentrations (27)□, and two bathypelagic (aphotic, 1250m and 1350m) samples.

We sought to describe the depth variations of prokaryotic community composition of Lake Baikal based on taxonomic profiles derived from metagenomes from winter and summer. Thus, read mapping was used to calculate the relative abundances of different phyla of Bacteria and Archaea across samples taken in winter (sub-ice) and summer water columns (Figure 1). As previously described, a clear distinction was observed between photic and aphotic samples (6)□. At all times, epipelagic samples tended to have higher abundances of Verrucomicrobiota, Actinobacteriota, Bacteroidota, and Cyanobacteria than their mesopelagic and bathypelagic counterparts. Acidobacteriota and Patescibacteria only displayed expressive abundances in the samples from the aphotic zone. Also, the abundances of Nitrospirota, Alphaproteobacteria, and Crenarchaeota increased towards the aphotic zone samples. Changes in energy availability brought by differences in light and temperature are major drivers of microbial community composition in aquatic habitats (28)□. The stable water temperatures in aphotic samples across seasons is likely responsible for the comparable community composition among these samples. Likewise, the more prominent change in temperature seen among photic zone samples was likely the driving factor behind the changes in prokaryotic community composition observed between seasons.

**Figure 1:**
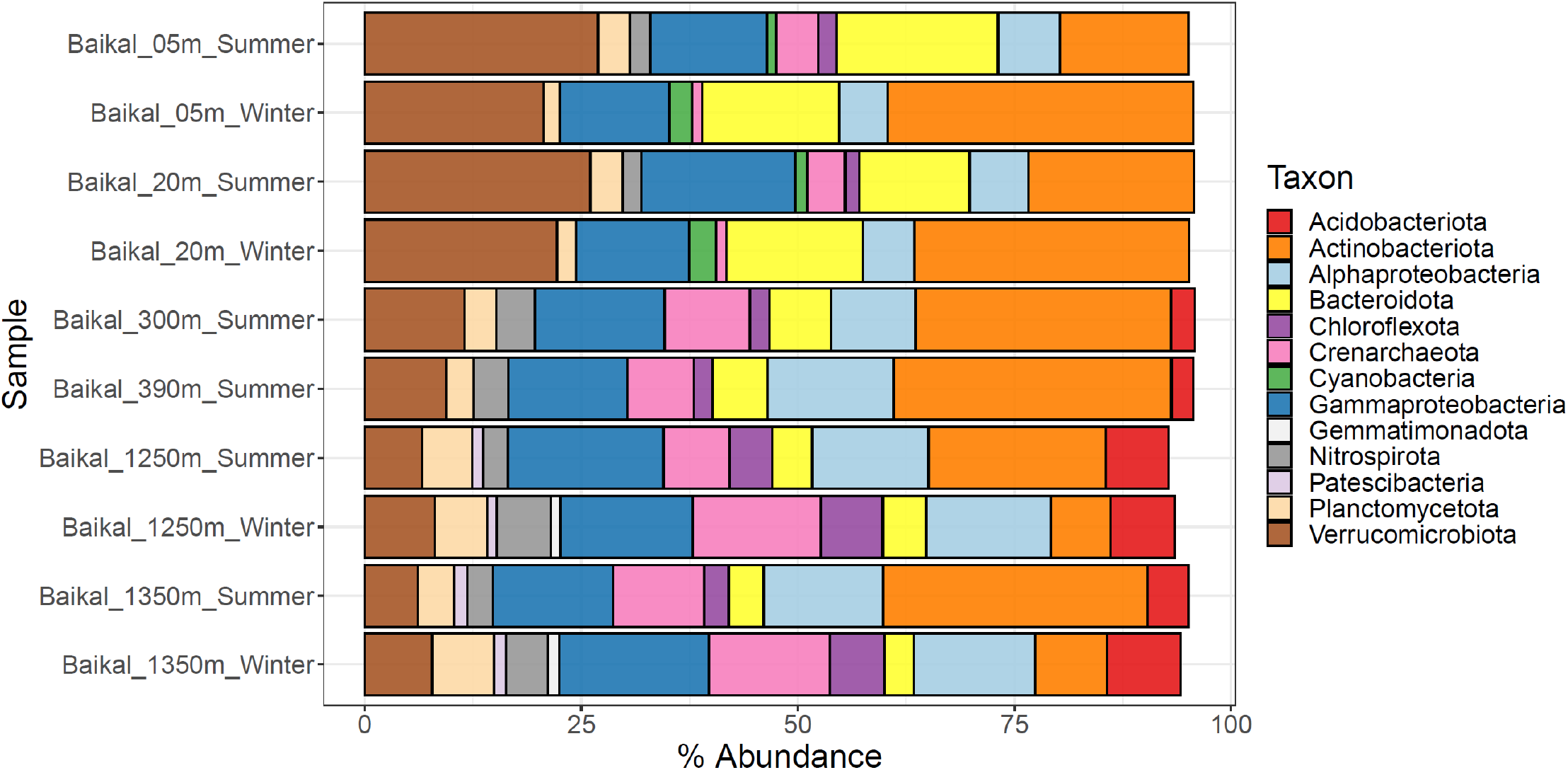
Lake Baikal prokaryotic community composition. Barplots depict the relative abundances of taxa of Archaea and Bacteria at the level of phylum (or class in the case of Proteobacteria) across the ten metagenomes from Lake Baikal. Only taxa that displayed relative abundances equal or above 1% are shown.

### Taxonomic classification and predicted hosts of Baikal viruses

The assembled scaffolds from ten Baikal metagenomes were analysed with VirSorter (29)□, VirFinder (30)□ and queried against the pVOGs database (31)□ to identify putative viral sequences. These putative viruses were subjected to manual curation after which a total of 19,475 sequences were classified as *bona fide* viruses (Table S1). Since these viral sequences were retrieved from metagenomes of the cellular fraction (as opposed to viromes), they are likely derived from viruses that were actively replicating at the time of sampling. The *bona fide* viral sequences were clustered into 9,916 viral populations on the basis of 95% average nucleotide identity and 80% shared genes within each population (19)□. Family level taxonomic assignments were achieved for 12,689 viral sequences. Most of them were classified into the families Myoviridae (7,155), Siphoviridae (3,138), Phycodnaviridae (1,195), and Podoviridae (809). The presence of viruses of eukaryotes in our dataset derives from the fact that samples were not pre-filtered to remove eukaryotic cells. Computational host prediction followed by manual curation allowed putative hosts at the taxonomic level of domain to be assigned to 2,870 viral sequences. These predictions suggested that the majority of these sequences belonged to viruses that infect Bacteria (2,135), but viruses that infect Archaea (29), Eukaryotes (621) and even virophages (85) were also identified. Among those assigned as viruses of bacteria (i.e. bacteriophages) the majority of sequences were predicted to infect Actinobacteria (640), followed by Proteobacteria (375), Bacteroidota (241) and Cyanobacteria (226). Although less frequent, some sequences were predicted to be derived from viruses that infect taxa with few or no isolated viruses such as Nitrospirota (9), Patescibacteria (14), and Crenarchaeota (23).

### Environmental drivers of viral community composition at Lake Baikal

We performed read recruitment from the 10 metagenomes to calculate relative abundances of viral sequences across samples. The resulting abundance matrix was used to investigate patterns of viral community composition in the Baikal ecosystem (Figure 2). These results pointed to a clear distinction between photic and aphotic samples regarding their viral community composition (Figure 2A). Among photic samples, a separation was observed between summer and winter samples which was mostly driven by viruses with high abundance among winter samples that displayed lower (sometimes below detection limit) abundances among the summer samples. No clear clustering of samples by season was observed among bathypelagic samples. All samples displayed comparable Shannon (8.0 – 9.1) and Simpson (0.9992 – 0.9996) diversity indexes, suggesting that despite the changes that take place in community composition across depth and seasons, the level of diversity within the communities remains stable.

**Figure 2:**
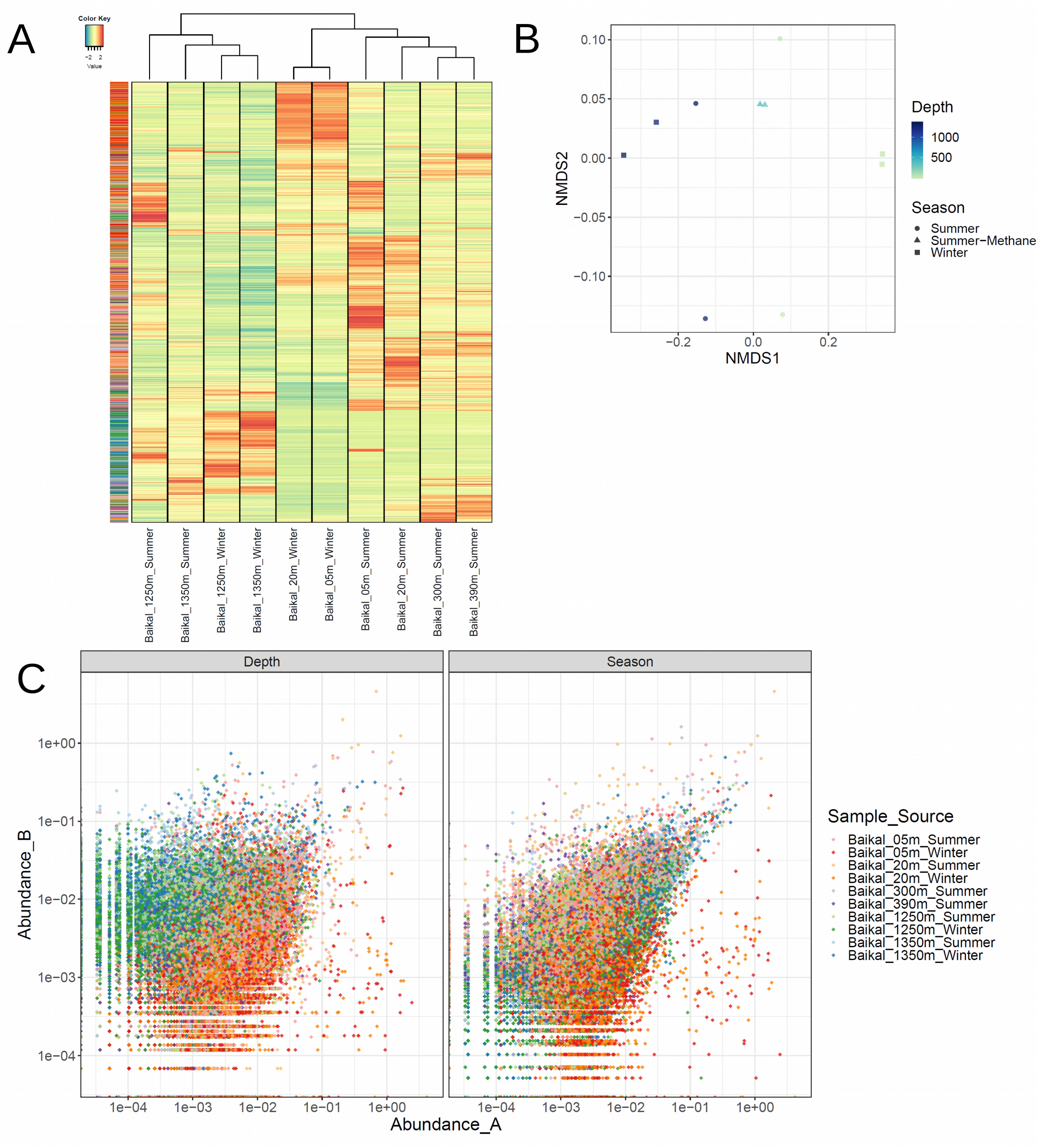
Lake Baikal viral community composition. A) Heatmap depicting the Z-score transformed abundances of 19,475 *bona fide* viral sequences across ten metagenomes from Lake Baikal. Both samples (columns) and viral sequences (rows) were subjected to hierarchical clustering based on Bray-Curtis dissimilarity distances. Side row colors indicate the sample from which each viral sequence was assembled. B) Non-metric multidimensional scaling comparison of the abundance of viral sequences across 10 baikal metagenomes based on Bray-Curtis dissimilarity distances. C) Scatterplots depicting the abundances of each viral scaffold paired by depth and season. In the left panel the relative abundances of sequences in the photic samples is displayed in the X axis while the abundance in the aphotic samples is displayed in the Y axis. Samples were paired as follows: 5m Winter x 1250m Winter; 5m Summer x 1250m Summer; 20m Winter x 1350m Winter; 20m Summer x 1350m Summer. In the right panel the relative abundances of sequences in the winter samples is displayed in the X axis while the abundance in the summer samples is displayed in the Y axis. Samples were paired as follows: 5m Winter x 5m Summer; 20m Winter x 20m Summer; 1250m Winter x 1250m Summer, 1350m Winter x 1350m Summer.

Non-metric multidimensional scaling also pointed to a clear distinction between samples from the photic and aphotic zones which were separated by NMDS1 (Figure 2B). However, no clear separation of samples by season was observed by NMDS1 or NMDS2. Next we analysed each individual scaffold by comparing abundances in the photic versus aphotic samples (from the same season), and also by comparing abundances in the summer versus winter samples (from the same depth). This result revealed the specific enrichment/depletion patterns of each viral sequence across the seasonal and bathymetric gradients (Figure 2C). Specifically, we observed distinctive clouds of viral sequences separating the photic from aphotic samples in the depth comparison, and the absence of a cloud separating winter from summer samples in the season comparison. This suggests that viruses specific of a given depth zone are much more frequent than viruses of a specific season.

Given these observations we next investigated how community composition changed according to the source, taxonomic affiliation, and predicted hosts of the viruses. Summing up the abundances of viral sequences according to the sample from which they were assembled revealed, on the one hand, that many of the viral sequences that were assembled from photic sample metagenomes were also abundant in the aphotic samples (Figure 3A). On the other hand, some viral sequences obtained from the aphotic samples were also abundant among the photic samples, albeit at lower relative abundances. Overall, this suggests an intense mixing between communities among zones, but with a greater influence of the photic zone over the aphotic zone, as could be expected from the convection currents in the lake (4,32)□. Next we summed up the abundances of viral sequences according to their family level taxonomic affiliation, obtained by closest relative assignment. This revealed a very stable trend of community composition with only very subtle changes in the relative abundances of the dominant families (Figure 3B). Overall, all samples were dominated by viruses assigned to the family Myoviridae, followed by Siphoviridae and Phycodnaviridae, with smaller contributions of Podoviridae and Mimiviridae. Finally, we summed up abundances of viral sequences according to the phylum of their assigned hosts. This pointed to more notable variations in community composition according to depth. Overall, the dominant groups in all samples were viruses predicted to infect Actinobacteriota and Proteobacteria (Figure 3C). The abundances of viruses predicted to infect Cyanobacteria decreased with depth, while the abundances of viruses predicted to infect Crenarchaeota, Chloroflexota, Planctomycetota, Nitrospirota, and Patescibacteria increased. Overall these results point to prominent changes in the composition of viral communities across the depth gradient, and subtle yet detectable differences across the seasonal changes. This is in agreement with recent findings that postulated that light and temperature are major drivers of viral community composition in marine ecosystems (19,33)□.

**Figure 3:**
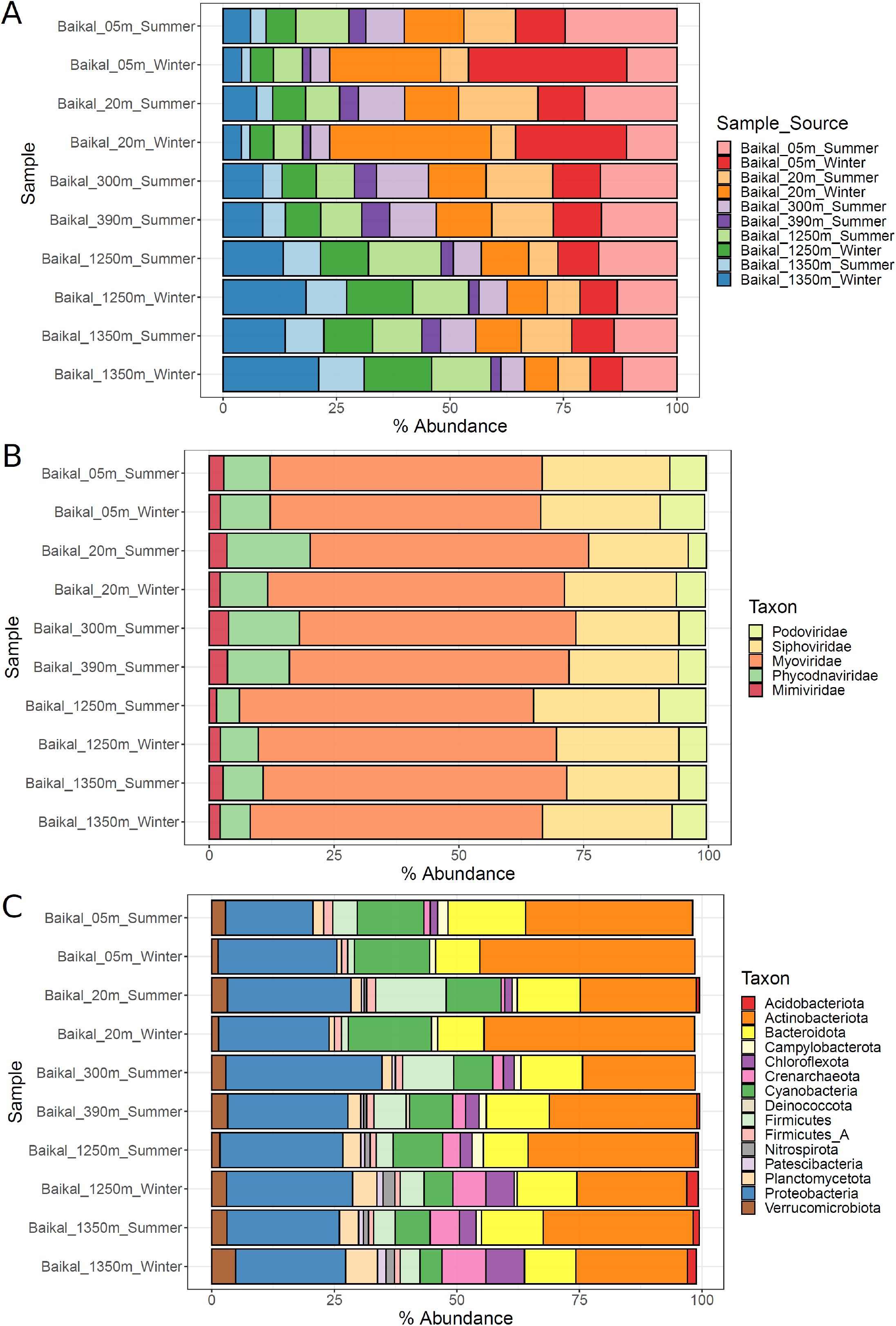
Barplots depicting the abundance of Baikal viruses summed up according to scaffold groups. A) Abundances summed up according to sample source of scaffolds B) Abundances summed up according to family level taxonomic classification of scaffolds. C) Abundances summed up according to predicted host phylum of scaffolds. Only families and host phyla that displayed abundances equal or above 0.5% are shown.

In previous studies we detected a predominance of freshwater microbes involved in the nitrification (i.e. Nitrospirota and Crenarchaeota) and oxidation of methyl compounds (i.e. Methylophilaceae) in the aphotic Lake Baikal (5,6)□. On the one hand, the ecological roles and diversity of AMGs of viruses that infect dominant groups of marine ecosystems (i.e. Cyanobacteria and Proteobacteria) has been characterized in detail (10,14,34)□. Likewise, the diversity of phages that infect Acnitobacteria (the dominant group among Baikal samples) in freshwater ecosystems has also been described in detail (20,21)□. On the other hand, the roles of viruses infecting nitrite oxidizers and methylotrophic bacteria in deep freshwater ecosystems is mostly unknown. Therefore, in this study we have focused on viruses that prey on microbes carrying out these processes, particularly viruses predicted to infect taxa for which few or no viruses have been described. In what follows we describe them and their potential involvement in biogeochemical processes through AMGs.

### Nitrospirota viruses from Lake Baikal interfere with dark carbon fixation

First, we manually curated the annotation of sequences of viruses predicted to infect bacteria of the phylum Nitrospirota. Members of Nitrospirota are chemolithoautrotrophic bacteria that perform nitrite oxidation mediated by nitrite oxidoreductases as a mean for energy acquisition, and some species are capable of complete nitrification (commamox) from ammonia to nitrate (35,36)□. These organisms use the reductive tricarboxylic acid (rTCA) cycle for dark carbon fixation (37,38)□. The viruses assigned to Nitrospirota were clustered into four distinct viral populations: VP_99, VP_1723, VP_4657 and VP_7454. Among those, there is considerable evidence suggesting that VP_99 (figure 4A) and VP_1723 (Figure 4B) are actual fragments of different regions of the same (or closely related) viral genome (Table S1). First, taxonomic classification assigned viruses from both populations to the genus T4Virus within the family Myoviridae. Second, the sequence representatives of both viral populations were assembled in the summer 1350m sample. Third, the representatives of these populations have almost identical GC content of 47.48% for VP_99 and 47.23% for VP_1723. Fourth, sequences from both populations match different regions of the Enterobacteria phage T4 genome (NC_000866.4). Finally, members of these two populations have a somewhat complementary gene content with the hallmark viral genes missing in one being present in the other.

**Figure 4:**
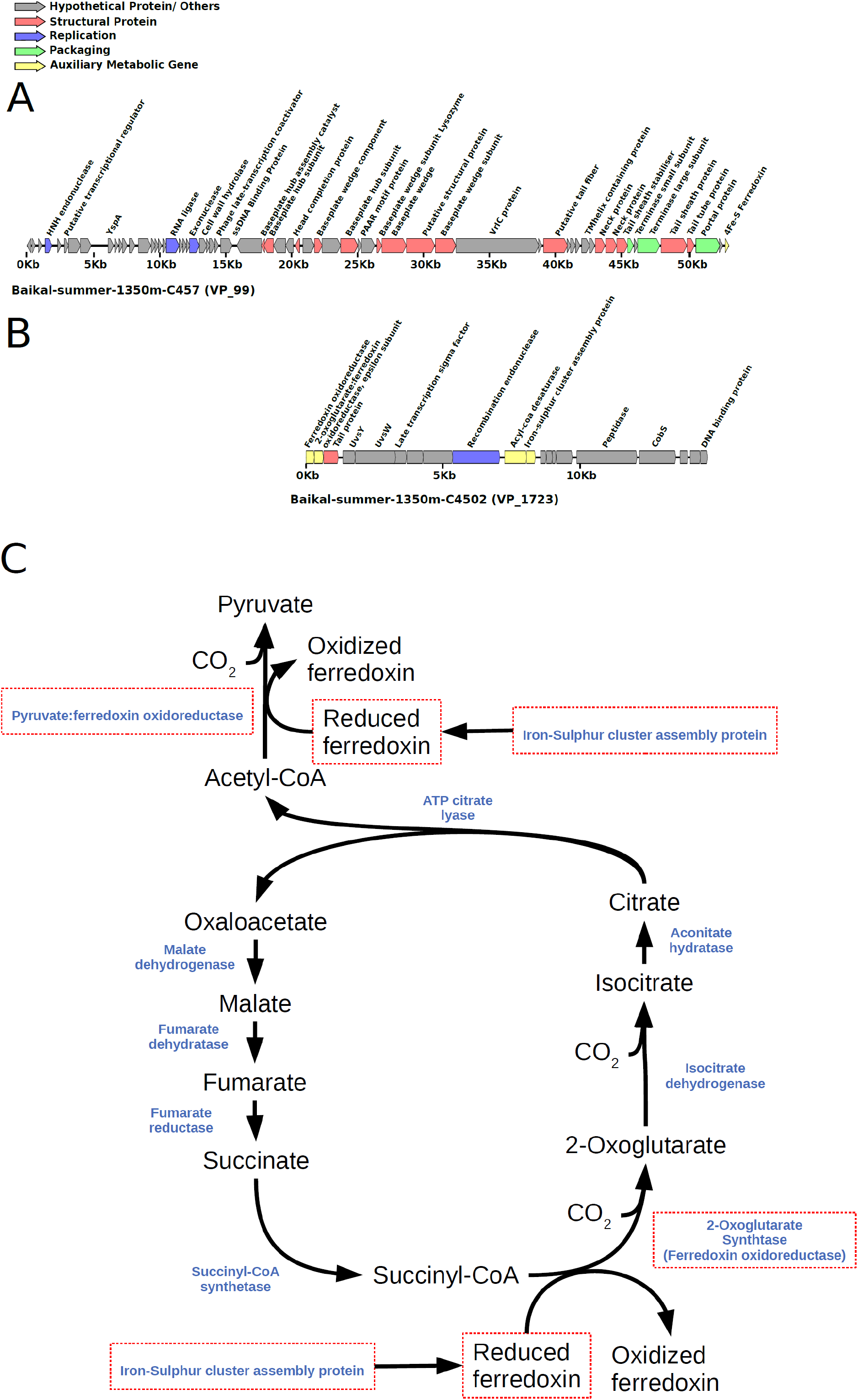
Novel viruses of Nitrospirota from Lake Baikal. A) Genomic map of Nitrospirota virus representative of VP_99. B) Genomic map of Nitrospirota virus representative of VP_1723. C) Reductive TCA cycle in Nitrospirota and potential influence of viruses over it. Enzymes are depicted in blue. Putative AMGs present in the genomes of either VP_99 or VP_1723 are highlighted by red rectangles.

The gene content of these populations provided insights into the infection strategies taken by these viruses (Figure 4). Most notably the members of VP_99 encoded a 4Fe-S Ferredoxin gene (Figure 4A). Ferredoxins are involved in a diverse set of redox reactions. These proteins are also involved in the energy metabolism of Nitrospirota (37)□ and on the rTCA cycle (38)□. The high degree of identity (86%) between viral and host ferredoxin suggests that this may be an AMG. Meanwhile, the members of VP_1723 displayed a different gene content (Figure 4B). Most notably, members of this population displayed a ferredoxin oxidoreductase, an epsilon subunit of 2-oxoglutarate:ferredoxin oxidoreductase, an Iron-Sulphur cluster biosynthesis protein, and an acyl-coA desaturase. All of these proteins had best hits to *Nitrospira* genes, suggesting that those are phage AMGs acquired from the host. The ferredoxin oxidoreductase and 2-oxoglutarate:ferredoxin oxidoreductase are clustered together in the genomes of *Nitrospira defluvii*, with the same orientation and little intergenic space, suggesting that they might have been acquired by the virus together in a single event and, more importantly, that they are all involved in the same cellular process.

The Iron-Sulphur cluster assembly protein is likely involved in the biosynthesis of the viral encoded ferredoxin (Figure 4C). Meanwhile, the 2-oxoglutarate:ferredoxin oxidoreductase is a key enzyme of the rTCA cycle in the genus *Nistrospira* (37,39)□. The viral ferredoxin oxidoreductase displayed significant homology with several ferredoxin oxidoreductases from the phylum Nitrospirota, including the pyruvate:ferredoxin oxidoreductase beta subunit of *Nitrospira defluvii*. This enzyme also mediates a key step of the reverse TCA cycle in *Nitrospira*. The presence of such genes in a viral genome is surprising since, to our knowledge, no AMGs acting on dark carbon fixation pathways have been described so far. Collectively, the occurrence of these genes in the viral genomes suggests that viruses of Nitrospirota modulate dark carbon fixation processes during infection. This is reminiscent to the way cyanophages modulate photosynthesis and carbon fixation pathways in Cyanobacteria (12)□.

The acyl-coA desaturase (also known as fatty acid desaturase or Stearoyl-CoA desaturase) is an enzyme that creates double bonds in fatty acids by removing hydrogen atoms, resulting in the creation of an unsaturated fatty acid. Unsaturated fatty acids are part of cell membranes, and a higher content of unsaturated fats is associated with higher membrane fluidity. The presence of an acyl-coa desaturase indicates that these viruses modulate the lipid metabolism of their host during infection. This gene belongs to a category of AMGs that is still poorly characterized in phages (14)□. Although eukaryotic viruses are known to influence the host lipid metabolism at multiple levels (40,41)□, a comprehensive understanding of this process in viruses of bacteria has not been achieved (42)□. Some ferredoxins are also involved in lipid metabolism (43)□, thus it is possible that the viral ferredoxins and acyl-coA desaturase work together to modulate host lipid metabolism during infection.

Based on these findings we postulate that Nitrospirota viruses of Lake Baikal make use of a diverse array of AMGs to modulate host metabolism during infection (Figure 4C). These findings have important implications to the understanding of dark carbon fixation in freshwater ecosystems, a process of recognized importance (44,45)□ in which the role of viruses is still poorly characterized. Our data demonstrates that viral enhanced dark carbon fixation is a process of ecological relevance. Specifically, our data suggests that viral mediated alterations to host metabolism could enhance the ratio of dark carbon fixation mediated by members of the phylum Nitrospirota, which account for up to 5 % of total microbes in bathypelagic waters of Lake Baikal. Thus, these viruses might play important roles in production of organic carbon by Nitrospirota that is eventually made available to the whole community following viral lysis. A previous publication reported the discovery of a Nistropirota virus from Lake Biwa, Japan (24)□. Nevertheless this sequence displayed no detectable homology to our viruses at the nucleotide level.

### Baikal viruses infecting methylotrophs interfere with methylotrophic metabolism and other major pathways

We identified viral populations predicted to infect methylotrophic bacteria. Among these, were included populations VP_139 (Figure 5A) and VP_266 (Figure 5B). These populations displayed ambiguous host predictions, with homology matches to multiple bacterial phyla (Bacteroidota and Proteobacteria). Hence, our original pipeline only assigned hosts to most members of these populations to the level of domain. Manual inspection of their computational host predictions revealed that homology matches to members of the family Methylophilaceae had higher bit-scores and identities and lower number of mismatches, indicating that members of VP_139 and VP_266 infect methylotrophic bacteria of the family Methylophilaceae, possibly from the closely related genera *Methylopumilus*, *Methylophilus* or *Methylotenera*. As before, we found evidence that these sequences are derived from the same genome (Table S1), as suggested by their complementary gene content, taxonomic affiliation (T4Virus), assembly source (winter surface samples), and GC content (35%).

Representative members of both VP_139 and VP_266 populations encoded methanol dehydrogenase (Figures 5A and 5B), a hallmark gene of methylotrophic metabolism in bacteria (46,47)□. This gene is responsible for the conversion of methanol into formaldehyde, the first and fundamental step of methylotrophic metabolism. To our knowledge, this is the first time this gene is being reported in viruses. We propose that viral methanol dehydrogenase is a novel AMG used by phages upon infection to boost up energy production of their methylotrophic hosts.

The representative sequence of VP_266 also encoded the pyrroloquinoline quinone precursor peptide PqqA. Pyrroloquinoline quinone (PQQ) is a redox cofactor which is necessary for the activity of the methanol dehydrogenase (48)□. The biosynthesis of PQQ is mediated by radical SAM proteins (49,50)□, which were detected in the genomes of VP_139. The representative sequence of VP_139 encoded also a methionine adenosyltransferase, which performs biosynthesis of S-adenosylmethionine (SAM) from L-methionine in the S-adenosyl-L-methionine cycle. In addition, it encoded an S-adenosyl-L-homocysteine hydrolase (5’-methylthioadenosine/S-adenosylhomocysteine nucleosidase) which performs the conversion of S-adenosyl-L-homocysteine into S-ribosyl-L-homocysteine also within this cycle. Methyltransferases, three of which were found in the representative genome of VP_139, also play a fundamental role on the S-adenosyl-L-methionine cycle, mediating the demethylation of S-adenosyl-L-methionine to convert it into S-adenosyl-L-homocysteine (51)□. The presence of so many auxiliary metabolic genes of the S-adenosyl-L-methionine cycle suggests that modulating this pathway is of fundamental relevance for the replication process of these viruses.

Members of VP_139 also encoded a phosphoribosylaminoimidazole synthetase (phosphoribosylformylglycinamide cyclo-ligase, *purM* gene), a widespread viral gene which is involved in nucleotide metabolism, and alpha and beta subunits for ribonucleotide-diphosphate reductase, which is also involved in this pathway. Together these observations suggest that members of VP_139 and VP_266 have a diverse array of proteins to modulate the metabolism of their methylotrophic hosts during infection (Figure 5C). This is achieved by expressing genes for methanol dehydrogenase and the cofactor PQQ to enhance the ratios of methanol oxidation to formaldehyde. It also expresses genes to boost up the biosynthesis of PQQ and the S-adenosyl-L-methionine cycle. Together these changes to host metabolism are likely to enhance the production of formaldehyde from methanol oxidation. The generated formaldehyde is then converted into formate through the tetrahydrofolate pathway or directed to the ribulose monophosphate cycle. The representative sequence of VP_266 encoded a peptide deformylase. This represents yet another candidate AMG, which would act to enhance the formate pool by removing formyl groups from host peptides. Interestingly, formate is used by phosphoribosylglycinamide formyltransferase 2 in the 5-aminoimidazole ribonucleotide biosynthesis pathway. Downstream of this step of the 5-aminoimidazole ribonucleotide biosynthesis pathway, phosphoribosylaminoimidazole synthetase and ribonucleotide-diphosphate reductase, that also participate in the biosynthesis of purines, were also found in the viral genomes. Thus, we conclude that these viruses enhance the methylotrophic metabolism of their hosts for the purpose of redirecting it towards the synthesis of nucleotides to be used in the replication of the viral genome (Figure 5C).

The discovery of these AMGs represents yet another novel way by which Baikal viruses modulate host metabolism. In this case it is of special relevance that these viruses affect three different host pathways: methanol oxidation, nucleotide metabolism, and the S-adenosyl-L-methionine cycle. In addition to this extensive gene repertoire we also identified other genes among these viral populations with the potential to be AMGs, albeit not directly linked to methanol oxidation or nucleotide metabolism. They included: glycerol-3-phosphate cytidylyltransferase which is involved in cell wall teichoic acid biosynthesis (52)□, and a class II aldolase/adducin family protein. Although our data does not allow us to determine the roles of these two proteins during infection, their presence among viral genomes is a novelty and it points to the diversity of strategies of these viruses to modulate host metabolism. A previous study has reported the isolation of a siphovirus (Phage P19250A) infecting *Methylopumilus planktonicus* (LD28) from Lake Soyang in South Korea. Nevertheless, this virus did not encode any of the putative AMGs reported here (53)□.

Another relevant microbe in the bathypelagic water column of Lake Baikal is *Methyloglobulus*, a genus of small (ca. 2.2 Mb of estimated genome size), yet abundant methanotrophs. These organisms were estimated to be among the most abundant microbes in bathypelagic waters of Lake Baikal (accounting up to 1 % of total mapped reads) and a MAG derived from this genus was described (6)□. We identified a viral population predicted to infect *Methyloglobulus*. In particular VP_1254 was composed of scaffolds of ca. 17 Kb that were assembled from bathypelagic metagenomes from both summer and winter (Figure 6A). These scaffolds displayed multiple homology matches to various taxa of Gammaproteobacteria. Among these, they consistently had high identity hits to a DnaK chaperone gene from the Baikal *Methyloglobulus* MAG. Finally, read recruitment confirmed the prevalence of these viruses among bathypelagic samples and absence from epipelagic and mesopelagic zones, following a pattern similar to that observed for *Methyloglobulus* (Figure. 6B). To our knowledge, no genomes of viruses infecting freshwater *Methyloglobulus* have been described. The gene content of these viruses included proteins involved in production of curly polymers and the ribosomal protein S21, which were previously detected in SAR11 phages (23)□ and a putative *Polynucleobacter* phage (5)□. Interestingly, these viruses did not encode the diverse array of AMGs described for the methylotroph viruses from VP_139 and VP_266, possibly because these sequences do not represent the complete viral genome. Another possible explanation is the fact that viruses from VP_139 and VP_266 are typical of the epipelagic zone, while those from VP_1254 are typical of the bathypelagic zone. Therefore, the metabolic constraints faced by these two groups of viruses during infection might be drastically different. These differences could explain the distinct array of AMGs between these two groups despite the fact that they infect closely related hosts with similar one-carbon metabolisms.

**Figure 5:**
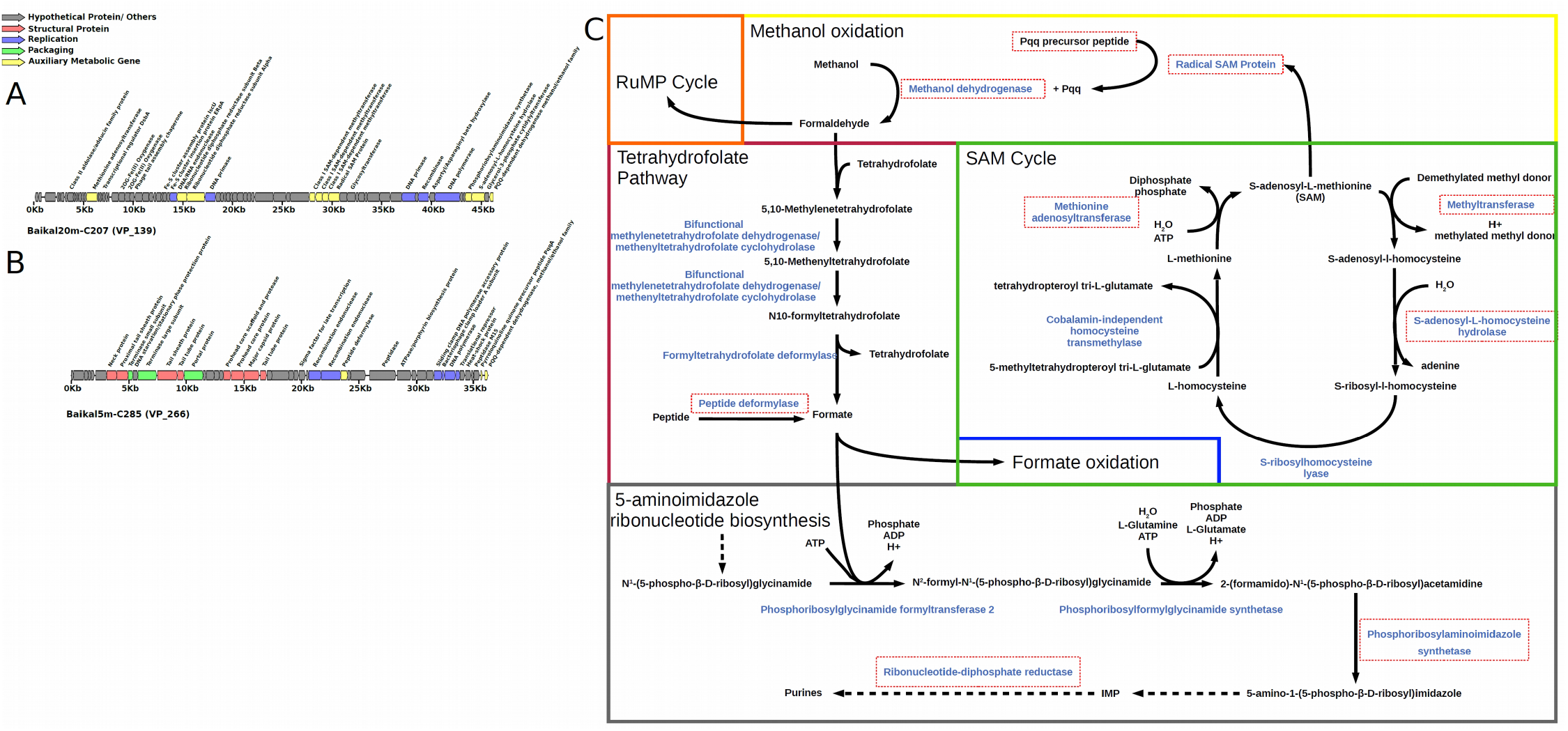
Novel viruses of Methylotrophs from Lake Baikal. A) Genomic map of virus representative of VP_139. B) Genomic map of virus representative of VP_266. C) Metabolic pathways of Methylotrophs and potential influence of viruses over it. Enzymes are depicted in blue. Putative AMGs present in the genomes of either VP_139 or VP_266 are highlighted by red rectangles. Colored rectangles separate different pathways/cycles. For simplicity, some reactions were omitted (represented by dashed arrows).

**Figure 6:**
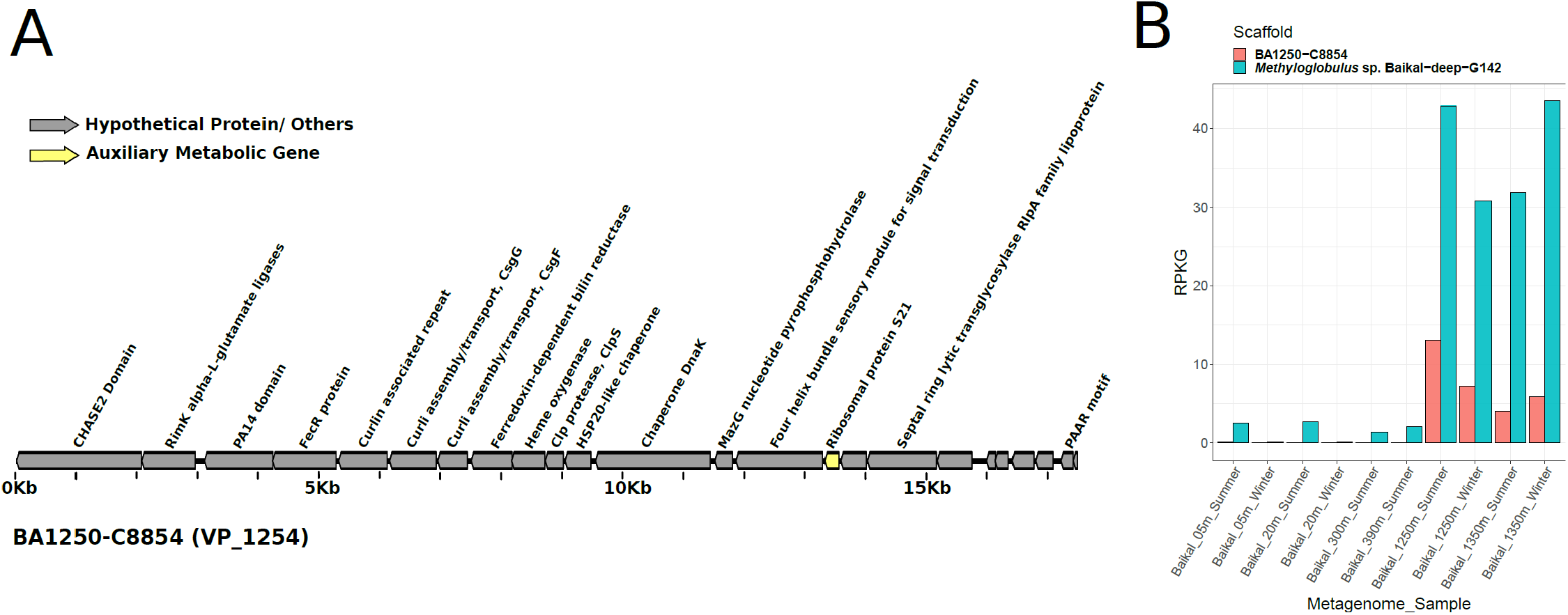
Novel *Methyloglobulus* virus from Lake Baikal. A) Genomic map of virus representative of VP_1254. B) Barplots depicting the abundances of the representative sequence of VP_1254 and its putative host MAG Methyloglobulus sp. Baikal-deep-G142 expressed as RPKG.

### Novel freshwater viruses of Crenarcheota

One of the singularities of the bathypelagic and mesopelagic Lake Baikal waters was the high abundances of Crenarchaeota (formerly Thaumarchaeota, e.g. *Nitrosopumilus* and *Nitrosoarchaeum*). We identified viral scaffolds predicted to infect Crenarchaeota in both summer and winter from bathypelagic and mesopelagic samples. Specifically, we retrieved scaffolds from multiple populations that presented a remarkable synteny to previously described marine Crenarchaeota viruses (Marthavirus) (54)□. The gene content of these scaffolds was conserved regardless of their sample of origin, as well as their gene order (Figure 7A). In addition, the typical Marthavirus genes radA, ATPases and CobS were conserved in their Lake Baikal counterparts. Thus, these viruses from Lake Baikal are the first representatives of freshwater viruses of Crenarchaeota, which are closely related to marine Marthavirus. However, a notorious difference between the marine and freshwater viruses of Crenarchaeota was the distribution of isoelectric points among their protein encoding genes (Figure 7B). Specifically, the isoelectric points of the Mediterranean Marthavirus representative was displaced towards more acidic values. This same tendency has been previously observed when comparing proteomes of Nitrosopumilaceae from marine and freshwater environments (55)□. This finding demonstrates that the shift in the distribution of isoeletric points among proteins that is observed during marine-freshwater transitions also extends to viruses, which sheds light on the processes by which these biological entities expand their ecological niches over time.

**Figure 7:**
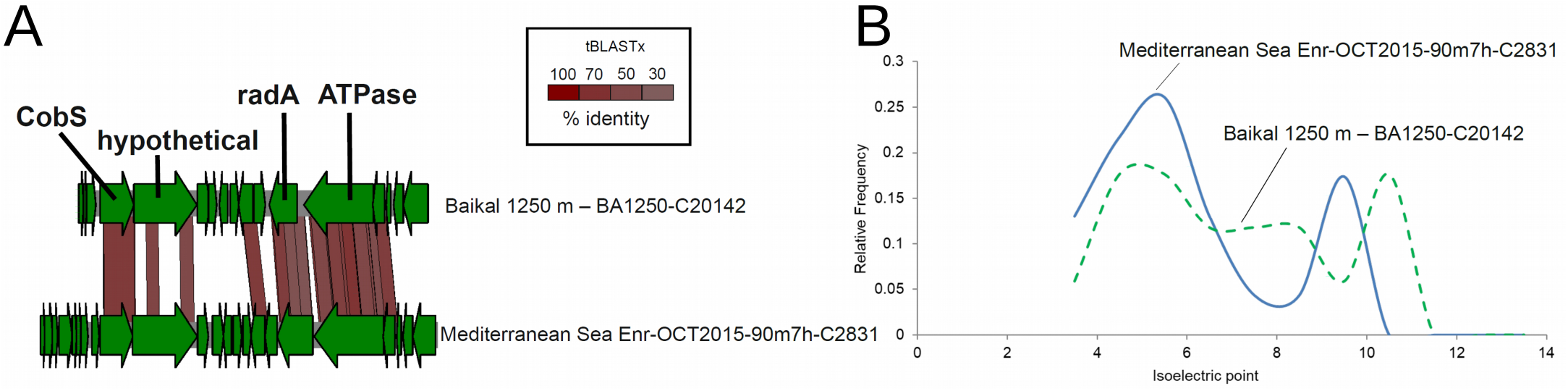
Novel Crenarchaeota viruses from Lake Baikal. A) Synteny maps depicting the similarities between a representative sequence of VP_2384 and a marine Marthavirus sequence. B) Distribution of isoeletric points among proteins from marine Marthaviruses and a close relative from Lake Baikal.

### Concluding remarks

We have taken advantage from the availability of metagenomes of Lake Baikal to shed light into the diversity of viruses within this unique habitat. Because we used metagenomes collected using a 0.2 μm filter we expected to obtain only the viral genomes that were being replicated within the retained cells. This method has been widely used and provides information about the viruses that are active in the community (20,56)□. Nevertheless, there might be other viruses that were missed by our approach but that might be recovered by sequencing the viral particles (virome). Even so we have obtained a large number of novel genomes of the most active viruses present in these samples. Interestingly, among them we have identified viruses predicted to prey on microbes that are major components of the community and which provide critical ecological functions. Specifically, in the large aphotic water mass of this deep lake.

We have found in Baikal another example of close relatives between marine and freshwater environments, the Crenarchaeota viruses. The degree of synteny observed between Marthaviruses and the Baikal scaffolds was remarkable. The occurrence of a large aerobic and deep water mass (a common feature between the ocean and Lake Baikal) is likely what facilitated the transition of these viruses between the two environments. Such parallelisms allow detection of specific adaptations required to live in low salt environments (the concentration of sodium for example is barely detectable in Baikal). One difference that has been detected in all cases of marine-freshwater transitions is the decrease in the isoelectric point of the proteome (55)□. The fact that this could also be detected in viruses indicates how critical this adaptation results for a proper functioning of basic molecular machinery such as that of DNA replication, transduction and translation.

In conclusion, our analysis of the viral communities from Lake Baikal has demonstrated how their composition and functioning changes across seasons and depths. These findings shed new light on the influence of environmental parameters over viruses in freshwater ecosystems. In addition, we described novel viruses with unique gene repertories, thus expanding the understanding of viral genetic diversity. These novel viruses also displayed new strategies for modulating host metabolism through auxiliary metabolic genes, by which they influence processes of ecological relevance, namely the methylotrophic metabolism and dark carbon fixation. Together these findings expand the understanding of viruses, the most abundant yet elusive biological entities on Earth and reveal novel roles played by them in processes of major biogeochemical relevance that take place in freshwater ecosystems.

## Methods

### Sampling and environmental parameters

The sampling strategy and sample post-processing for winter samples have been previously described (5,6)□. Summer samples were collected with the SBE 32 Carousel Water Sampler from aboard the RV ‘Vereshchagin’ in July 2018. Between 20 and 100 liters of water samples were retrieved from four horizons on each station. Water temperature and salinity were simultaneously measured with sensors SBE 19 Plus and SBE 25 Sealogger CTD (Sea-Bird Electronics) accurate within 0.002°C and with a resolution of 0.0003°C. pH values were measured using a pH 3310 meter (WTW, Germany). Overall, the hydrological conditions and the mineralization in the water column of the studied area corresponded to the data that were previously recorded during the same period in Lake Baikal (57,58)□. At Station 2, samples obtained on two runs were used to isolate DNA. The total volume of filtered water from the 300m sample was 70 L, and from the 390m sample the volume was 60 L.

For metagenomes, each sample was filtered through a net (size 27 μm) and then filtered through nitrocellulose filter with a pore size of 0.22 μm (Millipore, France), and the material from the filter was transferred to sterile flasks with 20 mL of lysis buffer (40 mmol L-1 EDTA, 50 mmol L-1 Tris/HCl, 0.75 mol L-1 sucrose) and stored at −20°C]. DNA was extracted according to the modified method of phenol-chloroformisoamyl alcohol method and stored at −70°C until further use. Metagenome sequencing, read-cleaning and assembly steps were performed as previously described (5,6)□.

### Sequence processing and analysis

Coding DNA sequences were identified in assembled scaffolds using Prodigal (59)□. Isoelectric points were calculated for each protein as previously described (55)□. Proteins sequences were queried against the NCBI-nr database using DIAMOND v0.8.22 (60)□ and Pfam using HMMER v3.1b2 (61)□ for taxonomic and functional annotation. Identification of putative viral sequences was performed in three steps: Sequences were analysed through VirSorter v1.0.6 (29)□ and those assigned to categories 1 and 2 and were considered as putative viruses. Also, sequences were analysed with VirFinder v1.1 (30)□ and those with a score ≥ 0.7 and p-value ≤ 0.05 were also considered putative viruses. Finally, protein sequences extracted from the scaffolds were queried against the pVOGs database (31)□ using HMMER set to a maximum e-value of 0.00001 (61)□. For each scaffold, we calculated the added viral quotient (AVQ) as the sum of the viral quotients of each pVOG that hits with the proteins of each scaffold (62)□. Scaffolds for which at least 20% of proteins mapped to pVOGs resulting in an AVQ ≥ 2 were considered putative viruses. Finally all of the putative viral sequences were subjected to manual inspection of their gene content and sequences that did not display a clearly viral signature (i.e. presence of hallmark viral genes and enrichment of hypothetical proteins) were excluded from further analysis resulting in a dataset of *bona fide* viral sequences. In addition, the *bona fide* viral sequences were clustered into viral populations based on 80% of shared genes at 95% average nucleotide identify as previously described (19)□.

### Taxonomic classification of viral sequences

Taxonomic affiliation of viral sequences was performed by closest relative affiliation. First, protein sequences derived from the *bona fide* viral sequences were queried against the viral sequences from the NCBI-nr database. DIAMOND was used with the following parameters: identity ≥ 30%, bit-score ≥ 50, alignment length ≥ 30 amino acids and e-value ≤ 0.00001 and the BLOSUM45 matrix. Next, the closest relative of each sequence was defined as the taxon that matched the highest number of protein sequences. Potential ties between taxa were resolved by selecting the one with the highest value of average identity among hits as the closest relative.

### Computational host prediction of viral sequences

Host predictions were performed based on previously reported benchmarking of methods to assign putative hosts to viruses based on shared genetic content between virus and host (63)□. For these searches, two reference databases were used: the NCBI RefSeq genomes of Bacteria and Archaea and a dataset of 266 prokaryote metagenome assembled genomes (MAGs) previously obtained from Lake Baikal (5,6)□. The taxonomic affiliation of RefSeq genomes was obtained from the Genomic Taxonomy Database (GTDB) (64)□. Baikal MAGs were also classified according to the GTDB system using GTDB-tk v0.3.2 (65)□. Three signals of virus-host association were analysed: homology matches, shared tRNAs, and CRISPR spacers. Homology matches were performed by querying viral sequences against the databases of prokaryote genomes using BLASTn v2.6.0+ (66)□. The cut-offs defined for these searches were: minimum alignment length of 300 bp, minimum identity of 50% and maximum e-value 0.001. tRNAs were identified in viral scaffolds using tRNAScan-SE v1.23 (67)□ using the bacterial models. The obtained viral tRNAs were queried against the database of prokaryote genomes using BLASTn. The cut-offs defined for these searches were: minimum alignment length of 60 bp, minimum identity of 90%, minimum query coverage of 95%, maximum of 10 mismatches and maximum e-value of 0.001. CRISPR spacers were identified in the databases of prokaryote genomes using CRISPRDetect v2.2 for the MAGs (68)□ and a custom script for the RefSeq genomes (69)□. The obtained spacers were queried against the sequences of *bona fide* viral sequences also using BLASTn. The cut-offs defined for these searches were: minimum identity of 95%, minimum query coverage of 95%, maximum of 1 mismatch and maximum e-value of 1. Ambiguous host predictions that assigned viruses to different microbial taxa were removed at each taxonomic level. Finally, putative hosts were also assigned to the *bona fide* viral sequences by manually inspecting their gene content.

### Prokaryote and viral abundance analysis

A database was compiled with one genome from each species representative of Bacteria and Archaea from the Genome Taxonomy Database (GTDB, release 89) (64)□. Protein sequences were predicted from these genomes using Prodigal v2.6.3 (59)□ with default parameters. Finally reads from the 10 metagenomes were queried against the GTDB database of protein encoding genes using DIAMOND (60)□ setting e-value to 0.00001 and minimum Bitscore to 50. For viruses, reads from the 10 metagenomes were queried against the assembled Baikal scaffolds using the sensitive-local mode of Bowtie2 v2.3.5.1 (70)□. The resulting abundance matrix was analysed using the Vegan Package (71)□ in R v3.6.1. Non-metric multidimensional scaling (NMDS) was performed based on the relative abundances of viral sequences using the Bray-Curtis dissimilarity measure.

## Data availability

Raw reads of winter (sub-ice) Lake Baikal metagenomes were previously published and are publicly available under the Bioproject numbers PRJNA396997 (SRR5896115 and SRR5896114 for 5 and 20 m samples, respectively) and PRJNA521725 (SRR8561390 and SRR8561391 for 1250 and 1350 m samples, respectively). Summer metagenomes have been deposited on NCBI SRA under bioproject number PRJNA615165. All assembled scaffolds were deposited at ENA under project number PRJEB37526.

## Code availability

All the relevant conde used in data analysis is publicly available.

## Acknowledgments

This work was supported by grants “VIREVO” CGL2016-76273-P [MCI/AEI/FEDER, EU] (cofounded with FEDER funds) from the Spanish Ministerio de Ciencia e Innovación and “HIDRAS3” PROMETEU/2019/009 from Generalitat Valenciana. FRV was also a beneficiary of the 5top100-program of the Ministry for Science and Education of Russia. FHC and PJCY were respectively supported by APOSTD/2018/186 and APOSTD/2019/009 post-doctoral fellowships from Generalitat Valenciana. RGS was supported by a predoctoral fellowship from the Valencian Consellería de Educació, Investigació, Cultura i Esport (ACIF/2016/050). The State Assignment 0345-2019-0007 supported the work (no. AAAA-A16-116122110064-7) of the Limnological Institute and grant OFIM no. 17-29-05040.

## Author contributions

FHC, PJCY, TIZ, ASZ, VGI and FRV conceived and designed experiments. PJCY, TIZ, ASZ, VGI and FRV collected samples and associated metadata. FHC, PJCY, RGS, RR and MLP analysed the data. All authors contributed to writing the manuscript.

## Ethics Approval and Consent to Participate

Not Applicable.

## Consent for publication

Not Applicable.

---

Table S1: Detailed description of all the analysed Baikal scaffolds. Fields include Completeness inferred by VirSorter. For each taxonomic level from domain to species: taxon name, number of CRISPR hits, number of homology matches hits, number of shared tRNA hits. Scaffold length. For each taxonomic level from domain to species: closest relative (CR) average amino acid identity (AAI), number of matched protein encoding genes (PEGs), percentage of matched PEGs relative to the total number of PEGs identified in the scaffold, and CR taxon name. MD5: MD5 checksum of scaffold sequence. Number of identified protein encoding genes. True_Virus: indicating if the scaffold was classified as a *bona fide* virus sequence. VP: Viral population to which scaffold was assigned. VirFinder score and p-value, VirSorter category, percentage of scaffold PEGs matched to pVOGs database, total number of hits to pVOGs database and added viral quotient (AVQ) of these hits.

## References

1. Kozhov MM. Biology of lake Baikal. Publ house Acad Sci USSR, Moscow. 1962;

2. Weiss RF, Carmack Carmack EC, Koropalov VM. Deep-water renewal and biological production in Lake Baikal. Nature [Internet]. 1991 Feb;349(6311):665–9. Available from: http://www.nature.com/articles/349665a0

3. Galazy GI. Atlas of Lake Baikal. GUGK, Moscow. 1993;489.

4. Shimaraev MN, Granin NG. On stratification and convection mechanism in Baikal. In: Dokl Akad Nauk SSSR. 1991. p. 381–5.

5. Cabello-Yeves PJ, Zemskay TI, Rosselli R, Coutinho FH, Zakharenko AS, Blinov VV, et al. Genomes of novel microbial lineages assembled from the sub-ice waters of Lake Baikal. Appl Environ Microbiol. 2018;84(1).

6. Cabello□Yeves PJ, Zemskaya TI, Zakharenko AS, Sakirko M V, Ivanov VG, Ghai R, et al. Microbiome of the deep Lake Baikal, a unique oxic bathypelagic habitat. Limnol Oceanogr [Internet]. 2019 Dec 30;lno.11401. Available from: https://onlinelibrary.wiley.com/doi/abs/10.1002/lno.11401

7. Suttle CA. Viruses in the sea. Nature [Internet]. 2005;437(7057):356–61. Available from: http://www.ncbi.nlm.nih.gov/pubmed/16163346

8. Danovaro R, Dell’Anno A, Corinaldesi C, Magagnini M, Noble R, Tamburini C, et al. Major viral impact on the functioning of benthic deep-sea ecosystems. Nature [Internet]. 2008 Aug;454(7208):1084–7. Available from: http://www.nature.com/articles/nature07268

9. Breitbart M. Marine Viruses: Truth or Dare. Ann Rev Mar Sci [Internet]. 2012 Jan 15;4(1):425–48. Available from: http://www.annualreviews.org/doi/10.1146/annurev-marine-120709-142805

10. Rosenwasser S, Ziv C, Creveld SG van, Vardi A. Virocell Metabolism: Metabolic Innovations During Host–Virus Interactions in the Ocean. Trends Microbiol [Internet]. 2016 Oct;24(10):821–32. Available from: http://linkinghub.elsevier.com/retrieve/pii/S0966842X16300695

11. Howard-Varona C, Lindback MM, Bastien GE, Solonenko N, Zayed AA, Jang H, et al. Phage-specific metabolic reprogramming of virocells. ISME J [Internet]. 2020 Jan 2; Available from: http://www.nature.com/articles/s41396-019-0580-z

12. Thompson LR, Zeng Q, Kelly L, Huang KH, Singer AU, Stubbe J, et al. Phage auxiliary metabolic genes and the redirection of cyanobacterial host carbon metabolism. Proc Natl Acad Sci [Internet]. 2011;108(39):E757–64. Available from: http://www.pnas.org/cgi/doi/10.1073/pnas.1102164108

13. Roux S, Hawley AK, Beltran MT, Scofeld M, Schwientek P, Stepanauskas R, et al. Ecology and evolution of viruses infecting uncultivated SUP05 bacteria as revealed by single-cell- and meta-genomics. Elife. 2014;2014(3):1–20.

14. Breitbart M, Bonnain C, Malki K, Sawaya NA. Phage puppet masters of the marine microbial realm. Nat Microbiol [Internet]. 2018 Jul 4;3(7):754–66. Available from: http://www.nature.com/articles/s41564-018-0166-y

15. Roux S, Brum JR, Dutilh BE, Sunagawa S, Duhaime MB, Loy A, et al. Ecogenomics and biogeochemical impacts of uncultivated globally abundant ocean viruses. Nature [Internet]. 2016;537(7622):589–693. Available from: http://biorxiv.org/content/early/2016/05/12/053090.abstract

16. Trubl G, Jang H Bin, Roux S, Emerson JB, Solonenko N, Vik DR, et al. Soil Viruses Are Underexplored Players in Ecosystem Carbon Processing. mSystems. 2018;3(5):1–21.

17. Coutinho FH, Silveira CB, Gregoracci GB, Thompson CC, Edwards RA, Brussaard CPD, et al. Marine viruses discovered via metagenomics shed light on viral strategies throughout the oceans. Nat Commun [Internet]. 2017 Dec 5;8(1):15955. Available from: http://dx.doi.org/10.1038/ncomms15955

18. Hurwitz BL, Hallam SJ, Sullivan MB. Metabolic reprogramming by viruses in the sunlit and dark ocean. Genome Biol [Internet]. 2013 Jan;14(11):R123. Available from: http://www.pubmedcentral.nih.gov/articlerender.fcgi?artid=4053976&tool=pmcentrez&rendertype=abstract

19. Brum JR, Ignacio-Espinoza JC, Roux S, Doulcier G, Acinas SG, Alberti A, et al. Patterns and ecological drivers of ocean viral communities. Science [Internet]. 2015 May 22 [cited 2015 May 23];348(6237):1261498. Available from: http://www.sciencemag.org/content/348/6237/1261498.short

20. Ghai R, Mehrshad M, Mizuno CM, Rodriguez-Valera F. Metagenomic recovery of phage genomes of uncultured freshwater actinobacteria. ISME J [Internet]. 2017 Jan 9;11(1):304–8. Available from: http://www.nature.com/articles/ismej2016110

21. Kavagutti VS, Andrei A-Ş, Mehrshad M, Salcher MM, Ghai R. Phage-centric ecological interactions in aquatic ecosystems revealed through ultra-deep metagenomics. Microbiome [Internet]. 2019 Dec 20;7(1):135. Available from: https://www.biorxiv.org/content/10.1101/670067v1

22. Chen L-X, Zhao Y-L, McMahon KD, Mori JF, Jessen GL, Nelson TC, et al. Wide distribution of phage that infect freshwater SAR11 bacteria. mSystems [Internet]. 2019;4(5):e00410–19. Available from: https://www.biorxiv.org/content/10.1101/672428v1

23. Zaragoza-Solas A, Rodriguez-Valera F, López-Pérez M. Metagenome Mining Reveals Hidden Genomic Diversity of Pelagimyophages in Aquatic Environments. mSystems [Internet]. 2020;5(1):1–16. Available from: http://msystems.asm.org/lookup/doi/10.1128/mSystems.00905-19

24. Okazaki Y, Nishimura Y, Yoshida T, Ogata H, Nakano S. Genome resolved viral and cellular metagenomes revealed potential key virus host interactions in a deep freshwater lake. Environ Microbiol [Internet]. 2019 Dec 6;21(12):4740–54. Available from: https://onlinelibrary.wiley.com/doi/abs/10.1111/1462-2920.14816

25. Butina TV, Bukin YS, Khanaev IV, Kravtsova LS, Maikova OO, Tupikin AE, et al. Metagenomic analysis of viral communities in diseased Baikal sponge Lubomirskia baikalensis. Limnol Freshw Biol. 2019;2019(1):155–62.

26. Potapov SA, Tikhonova I V, Krasnopeev AY, Kabilov MR, Tupikin AE, Chebunina NS, et al. Metagenomic Analysis of Virioplankton from the Pelagic Zone of Lake Baikal. Viruses [Internet]. 2019 Oct 29;11(11):991. Available from: https://www.mdpi.com/1999-4915/11/11/991

27. Granin NG, Makarov MM, Kucher KM, Gnatovsky RY. Gas seeps in Lake Baikal—detection, distribution, and implications for water column mixing. Geo-Marine Lett. 2010;30(3–4):399–409.

28. Sunagawa S, Coelho LP, Chaffron S, Kultima JR, Labadie K, Salazar G, et al. Structure and function of the global ocean microbiome. Science (80-) [Internet]. 2015 May 22;348(6237):1261359–1261359. Available from: http://www.sciencemag.org/cgi/doi/10.1126/science.1261359

29. Roux S, Enault F, Hurwitz BL, Sullivan MB. VirSorter: mining viral signal from microbial genomic data. PeerJ. 2015;3:e985.

30. Ren J, Ahlgren NA, Lu YY, Fuhrman JA, Sun F. VirFinder: a novel k-mer based tool for identifying viral sequences from assembled metagenomic data. Microbiome [Internet]. 2017;5(1):69. Available from: http://microbiomejournal.biomedcentral.com/articles/10.1186/s40168-017-0283-5

31. Grazziotin AL, Koonin E V., Kristensen DM. Prokaryotic Virus Orthologous Groups (pVOGs): a resource for comparative genomics and protein family annotation. Nucleic Acids Res [Internet]. 2017 Jan 4;45(D1):D491–8. Available from: https://academic.oup.com/nar/article-lookup/doi/10.1093/nar/gkw975

32. Shimaraev MN, Granin NG, Gnatovskij RJ, Blinov V V. The mechanism of oxygen aeration of bottom waters of Lake Baikal. Dokl Earth Sci. 2015;461(2):379–83.

33. Hurwitz BL, Brum JR, Sullivan MB. Depth-stratified functional and taxonomic niche specialization in the ‘core’ and ‘flexible’ Pacific Ocean Virome. ISME J [Internet]. 2015;9:472–84. Available from: http://www.nature.com/doifinder/10.1038/ismej.2014.143

34. Hurwitz BL, U’Ren JM. Viral metabolic reprogramming in marine ecosystems. Curr Opin Microbiol [Internet]. 2016;31:161–8. Available from: http://www.sciencedirect.com/science/article/pii/S1369527416300376

35. van Kessel MAHJ, Speth DR, Albertsen M, Nielsen PH, Op den Camp HJM, Kartal B, et al. Complete nitrification by a single microorganism. Nature [Internet]. 2015;528(7583):555–9. Available from: http://www.nature.com/doifinder/10.1038/nature16459

36. Daims H, Lebedeva EV, Pjevac P, Han P, Herbold C, Albertsen M, et al. Complete nitrification by Nitrospira bacteria. Nature [Internet]. 2015;528(7583):504–9. Available from: http://www.nature.com/doifinder/10.1038/nature16461%5Cnhttp://dx.doi.org/10.1038/nature16461

37. Lücker S, Wagner M, Maixner F, Pelletier E, Koch H, Vacherie B, et al. A Nitrospira metagenome illuminates the physiology and evolution of globally important nitrite-oxidizing bacteria. Proc Natl Acad Sci U S A. 2010;107(30):13479–84.

38. Hügler M, Sievert SM. Beyond the Calvin Cycle: Autotrophic Carbon Fixation in the Ocean. Ann Rev Mar Sci [Internet]. 2011 Jan 15;3(1):261–89. Available from: http://www.ncbi.nlm.nih.gov/pubmed/21329206

39. Daims H, Lücker S, Wagner M. A New Perspective on Microbes Formerly Known as Nitrite-Oxidizing Bacteria. Trends Microbiol [Internet]. 2016;24(9):699–712. Available from: http://dx.doi.org/10.1016/j.tim.2016.05.004

40. Heaton NS, Randall G. Multifaceted roles for lipids in viral infection. Trends Microbiol [Internet]. 2011;19(7):368–75. Available from: http://dx.doi.org/10.1016/j.tim.2011.03.007

41. Lange PT, Lagunoff M, Tarakanova VL. Chewing the fat: The conserved ability of DNA viruses to hijack cellular lipid metabolism. Viruses. 2019;11(2):1–19.

42. Kutter E, Bryan D, Ray G, Brewster E, Blasdel B, Guttman B. From host to phage metabolism: Hot tales of phage T4’s takeover of E. coli. Viruses. 2018;10(7).

43. Yang W, Wittkopp TM, Li X, Warakanont J, Dubini A, Catalanotti C, et al. Critical role of chlamydomonas reinhardtii ferredoxin-5 in maintaining membrane structure and dark metabolism. Proc Natl Acad Sci U S A. 2015 Dec 1;112(48):14978–83.

44. Casamayor EO, García-Cantizano J, Pedrós-Alió C. Carbon dioxide fixation in the dark by photosynthetic bacteria in sulfide-rich stratified lakes with oxic-anoxic interfaces. Limnol Oceanogr [Internet]. 2008 Jul 1 [cited 2020 Feb 11];53(4):1193–203. Available from: http://doi.wiley.com/10.4319/lo.2008.53.4.1193

45. Santoro AL, Bastviken D, Gudasz C, Tranvik L, Enrich-Prast A. Dark Carbon Fixation: An Important Process in Lake Sediments. PLoS One. 2013;8(6):1–7.

46. Chistoserdova L. Modularity of methylotrophy, revisited. Environ Microbiol. 2011;13(10):2603–22.

47. Salcher MM, Schaefle D, Kaspar M, Neuenschwander SM, Ghai R. Evolution in action: habitat transition from sediment to the pelagial leads to genome streamlining in Methylophilaceae. ISME J [Internet]. 2019 Nov 10;13(11):2764–77. Available from: http://www.nature.com/articles/s41396-019-0471-3

48. Anthony C, Williams P. The structure and mechanism of methanol dehydrogenase. Biochim Biophys Acta - Proteins Proteomics. 2003;1647(1–2):18–23.

49. Wecksler SR, Stoll S, Tran H, Magnusson OT, Wu SP, King D, et al. Pyrroloquinoline quinone biogenesis: Demonstration that PqqE from Klebsiella pneumoniae is a radical S-adenosyl-L-methionine enzyme. Biochemistry. 2009;48(42):10151–61.

50. Barr I, Latham JA, Iavarone AT, Chantarojsiri T, Hwang JD, Klinman JP. Demonstration that the radical s-adenosylmethionine (SAM) Enzyme PqqE catalyzes de novo carbon-carbon cross-linking within a peptide substrate PqqA in the presence of the peptide chaperone PqqD. J Biol Chem. 2016;291(17):8877–84.

51. Fontecave M, Atta M, Mulliez E. S-adenosylmethionine: Nothing goes to waste. Trends Biochem Sci. 2004;29(5):243–9.

52. Young Seo Park, Sweitzer TD, Dixon JE, Kent C. Expression, purification, and characterization of CTP:glycerol-3-phosphate cytidylyltransferase from Bacillus subtilis. J Biol Chem. 1993;268(22):16648–54.

53. Moon K, Kang I, Kim S, Kim SJ, Cho JC. Genome characteristics and environmental distribution of the first phage that infects the LD28 clade, a freshwater methylotrophic bacterial group. Environ Microbiol. 2017;19(11):4714–27.

54. López□Pérez M, Haro□Moreno JM, de la Torre JR, Rodrigue□Valera F. Novel Caudovirales associated with Marine Group I Thaumarchaeota assembled from metagenomes. Environ Microbiol [Internet]. 2019 Jun 3;21(6):1980–8. Available from: http://doi.wiley.com/10.1111/1462-2920.14462

55. Cabello-Yeves PJ, Rodriguez-Valera F. Marine-freshwater prokaryotic transitions require extensive changes in the predicted proteome. Microbiome. 2019;7(1):1–12.

56. López-Pérez M, Haro-Moreno JM, Gonzalez-Serrano R, Parras-Moltó M, Rodriguez-Valera F. Genome diversity of marine phages recovered from Mediterranean metagenomes: Size matters. PLoS Genet. 2017;13(9):e1007018.

57. Votintsev KK, Meshcheryakova AI, Popovskaya GI. Cycle of organic matter in Lake Baikal. Novosib Nauk. 1975;

58. Khodzher T, Domysheva VM, Sorokovikova LM, Golobokova LP. Methods for monitoring the chemical composition of Lake Baikal water. In: Novel methods for monitoring and managing land and water resources in Siberia. Springer; 2016. p. 113–32.

59. Hyatt D, Chen G-L, Locascio PF, Land ML, Larimer FW, Hauser LJ. Prodigal: prokaryotic gene recognition and translation initiation site identification. BMC Bioinformatics [Internet]. 2010;11:119. Available from: http://www.pubmedcentral.nih.gov/articlerender.fcgi?artid=2848648&tool=pmcentrez&rendertype=abstract

60. Buchfink B, Xie C, Huson DH. Fast and sensitive protein alignment using DIAMOND. Nat Methods [Internet]. 2015 Jan 17;12(1):59–60. Available from: http://www.nature.com/articles/nmeth.3176

61. Finn RD, Clements J, Arndt W, Miller BL, Wheeler TJ, Schreiber F, et al. HMMER web server: 2015 update. Nucleic Acids Res. 2015;43(W1):W30--W38.

62. Coutinho FH, Edwards RA, Rodríguez-Valera F. Charting the diversity of uncultured viruses of Archaea and Bacteria. BMC Biol [Internet]. 2019 Dec 29;17(1):109. Available from: https://www.biorxiv.org/content/10.1101/480491v1.full

63. Edwards RA, McNair K, Faust K, Raes J, Dutilh BE. Computational approaches to predict bacteriophage–host relationships. Smith M, editor. FEMS Microbiol Rev [Internet]. 2016 Mar;40(2):258–72. Available from: https://academic.oup.com/femsre/article-lookup/doi/10.1093/femsre/fuv048

64. Parks DH, Chuvochina M, Waite DW, Rinke C, Skarshewski A, Chaumeil P-A, et al. A standardized bacterial taxonomy based on genome phylogeny substantially revises the tree of life. Nat Biotechnol [Internet]. 2018 Nov 27;36(10):996–1004. Available from: https://www.biorxiv.org/content/early/2018/01/30/256800

65. Chaumeil P-A, Mussig AJ, Hugenholtz P, Parks DH. GTDB-Tk: a toolkit to classify genomes with the Genome Taxonomy Database. Hancock J, editor. Bioinformatics [Internet]. 2019 Nov 15 [cited 2020 Jan 11]; Available from: https://academic.oup.com/bioinformatics/advance-article/doi/10.1093/bioinformatics/btz848/5626182

66. Altschul SF, Gish W, Miller W, Myers EW, Lipman DJ. Basic local alignment search tool. J Mol Biol [Internet]. 1990;215(3):403–10. Available from: http://www.ncbi.nlm.nih.gov/pubmed/2231712

67. Lowe TM, Chan PP. tRNAscan-SE On-line: integrating search and context for analysis of transfer RNA genes. Nucleic Acids Res [Internet]. 2016 Jul 8 [cited 2020 Jan 16];44(W1):W54–7. Available from: https://academic.oup.com/nar/article-lookup/doi/10.1093/nar/gkw413

68. Biswas A, Staals RHJ, Morales SE, Fineran PC, Brown CM. CRISPRDetect: A flexible algorithm to define CRISPR arrays. BMC Genomics [Internet]. 2016 Dec 17 [cited 2020 Jan 16];17(1):356. Available from: http://bmcgenomics.biomedcentral.com/articles/10.1186/s12864-016-2627-0

69. Díez-Villaseñor C, Rodriguez-Valera F. CRISPR analysis suggests that small circular single-stranded DNA smacoviruses infect Archaea instead of humans. Nat Commun [Internet]. 2019;10(1):294. Available from: http://www.nature.com/articles/s41467-018-08167-w

70. Langmead B, Salzberg SL. Fast gapped-read alignment with Bowtie 2. Nat Methods [Internet]. 2012 Apr 4 [cited 2014 Jul 10];9(4):357–9. Available from: http://dx.doi.org/10.1038/nmeth.1923

71. Oksanen J. Vegan: an introduction to ordination. Management [Internet]. 2008;1:1–10. Available from: http://doi.acm.org/10.1145/2037556.2037605%5Cnftp://ftp3.ie.freebsd.org/pub/cran.r-project.org/web/packages/vegan/vignettes/intro-vegan.pdf

